# Transcriptome network analysis implicates CX3CR1-positive type 3 dendritic cells in non-infectious uveitis

**DOI:** 10.1101/2021.11.16.468816

**Authors:** S. Hiddingh, A. Pandit, F.H. Verhagen, R. Rijken, N. H. Servaas, C.G.K. Wichers, N.H. ten Dam-van Loon, S.M. Imhof, T.R.D.J. Radstake, J.H. de Boer, J.J.W. Kuiper

## Abstract

**Background:** Type I interferons (IFNs) promote the expansion of subsets of CD1c+ conventional dendritic cells (CD1c+ DCs), but the molecular basis of CD1c+ DCs involvement in conditions not associated without elevated type I IFNs remains unclear.

**Methods:** We analyzed CD1c+ DCs from two cohorts of non-infectious uveitis patients and healthy donors using RNA-sequencing followed by high-dimensional flow cytometry to characterize the CD1c+ DC populations.

**Results:** We report that the CD1c+ DCs pool from patients with non-infectious uveitis is skewed towards a gene module with the chemokine receptor *CX3CR1* as the key hub gene. We confirmed these results in an independent case-control cohort and show that the disease-associated gene module is not mediated by type I IFNs. An analysis of peripheral blood using flow cytometry revealed that CX3CR1+ DC3s were diminished, whereas CX3CR1-DC3s were not. Stimulated CX3CR1+ DC3s secrete high levels of inflammatory cytokines, including TNF-alpha, and CX3CR1+ DC3-like cells can be detected in inflamed eyes of patients.

**Conclusions:** These results show that CX3CR1+ DC3s are implicated in non-infectious uveitis and can secrete proinflammatory mediators implicated in its pathophysiology.

**Funding:** The presented work is supported by UitZicht (project number #2014-4, #2019-10, an #2021-4). The funders had no role in the design, execution, interpretation, or writing of the study.

## INTRODUCTION

Non-infectious uveitis refers to a group of chronic inflammatory eye diseases that are among the leading causes of preventable vision loss in western countries (*1,2*). Currently, little is known about the disease mechanism of non-infectious uveitis. A large body of mechanistic studies using experimental autoimmune uveitis (EAU) in rodents suggest that T cells play a role in non-infectious uveitis (3,4). Human non-infectious uveitis is characterized by inflammation and T cells infiltrating the eyes through unknown mechanisms. The genetic association between non-infectious uveitis and *MHC*, and *ERAP1, ERAP2* genes indicates that antigen presentation is central to the etiology (*5–8*). A key antigen presenting cell type are dendritic cells. Despite their important role in EAU, dendritic cells have yet to be fully investigated in human non-infectious uveitis (9–11). CD1c-positive conventional dendritic cells (CD1c+ DCs) have been found to be associated with disease activity (12–14), and are abundant in eye fluid of patients (*15*). In order to understand the role of CD1c+ DCs in non-infectious uveitis, it is necessary to understand their functions.

Several single-cell studies have revealed that the CD1c+ DCs (and its murine equivalent, termed “cDC2s”) consists of multiple subsets derived from distinct progenitors (*16–18*). The type I Interferon (IFN) family of cytokines promotes the expansion of a subset of CD1c+ DCs called “DC3” (*17,19,20*). DC3s are increased in blood of type I IFN-driven *systemic lupus erythematosus* (SLE) patients (17). A significant difference between non-infectious uveitis and SLE is that active uveitis is accompanied by lower levels of type I IFN (*21,, 22*). It should be noted that although type I interferons can induce lupus-like disease, they can also suppress non-infectious uveitis (*21,23,24*), pointing to an alternative disease mechanism implicating CD1c+ DCs in non-infectious uveitis. Hence, we do not fully understand the characteristics of CD1c+ DC during autoimmunity, especially in conditions not driven by type I IFNs.

For the purpose of characterizing the core transcriptional features and subset composition of CD1c+ DCs in autoimmunity of the eye, we used whole transcriptome profiling by bulk RNA-sequencing of peripheral blood CD1c+ DCs and multiparameter flow cytometry of two cohorts of non-infectious uveitis patients and healthy donors. We constructed co-expression networks that identified a robust gene module associated with non-infectious uveitis in patients that helped identify a CX3CR1-positive CD1c+ DC subset.

## MATERIAL AND METHODS

### Patients and patient material

This study was conducted in compliance with the Helsinki principles. Ethical approval was requested and obtained from the Medical Ethical Research Committee in Utrecht (METC protocol number #14-065/M). All patients signed written informed consent before participation. We collected blood from a discovery cohort of 29 and a replication cohort of 22 adult patients (**Table 1**) with HLA-B27-associated Acute Anterior Uveitis (AU), Idiopathic Intermediate Uveitis (IU), or HLA-A29-associated Birdshot Uveitis (BU). Patients were recruited at the outbound patient clinic of the department of Ophthalmology of the University Medical Center Utrecht between July 2014 and January 2017. We recruited twenty-nine age and sex matched anonymous blood donors of European Ancestry with no history of ocular inflammatory disease at the same institute to serve as unaffected controls (**Table 1**). Uveitis was classified and graded in accordance with the SUN classification (25). Each patient underwent a full ophthalmological examination by an ophthalmologist experienced in uveitis, routine laboratory screening, and an X-Ray of the lungs. Laboratory screening included erythrocyte sedimentation rate, renal and liver function tests, angiotensin converting enzyme (ACE), and screening for infectious agents in the serum and an Interferon-Gamma Release Assay (IGRA) was obtained for all patients. All patients with AU and BU were HLA-B27 or HLA-A29-positive, respectively (confirmed by HLA typing). All patients had active uveitis (new onset or relapse) and there was no clinical evidence for uveitis-associated systemic inflammatory disease (e.g., rheumatic condition) till the time of sampling. None of the patients received systemic immunomodulatory treatment in the last 3 months, other than low dose (≤10mg) oral prednisolone in one BU patient of cohort II and one AU patient of cohort I.

**Table 1.** Characteristics of the patients and controls from cohort I and cohort II. Abbreviations: BU: Birdshot uveitis, AU: HLA-B27 associated anterior uveitis, HC: healthy control, IU: idiopathic intermediate uveitis, n.a.: not applicable, * Fisher’s exact test, ** ANOVA,*** Kruskal-Wallis.

| Cohort I | AU | IU | BU | HC | *P* value |
| --- | --- | --- | --- | --- | --- |
| **N** | 10 | 5 | 8 | 13 | Total=36 |
| **Male / Female** | 2/8 | 3/2 | 5/3 | 5/8 | 0.26* |
| **Age in years; mean ± SD** | 45 ± 16 | 30 ± 9 | 42 ± 10 | 42 ± 13 | 0.24*** |
| **Disease duration in years; median (range)** | 8.1 (0.2-22.3) | 3.4 (0.4-14.1) | 0.9 (0.2-19.9) | n.a. | 0.36** |
| Cohort II | AU | IU | BU | HC | *P* value |
| **N** | 9 | 9 | 10 | 14 | Total=42 |
| **Male / Female** | 3/6 | 2/7 | 4/6 | 6/8 | 0.8790* |
| **Age in years; mean ± SD** | 47. ± 17 | 39 ± 14 | 52 ± 13 | 39 ± 10 | 0.06*** |
| **Disease duration in years; median (range)** | 5.8 (0.1-39.3) | 3.7 (0.2-20.0) | 1.3 (0.2-15.1) | n.a. | 0.14** |

### CD1c+ DC purification

Peripheral blood mononuclear cells (PBMCs) were isolated by standard ficoll density gradient centrifugation from 70mL heparinized blood immediately after blood withdrawal (GE Healthcare, Uppsala, Sweden). For the first cohort, ten batches (individual days) of 4-5 randomly selected patient and control samples of nitrogen stored PBMCs (mean storage time of 11 [range 0-31] months) were carefully thawed and subjected to sorting by the BD FACSAria™ III sorter after incubation with a panel of surface antibodies (**Supplementary File 1A**) and FACS buffer (1% bovine serum albumin and 0.1% sodium azide in phosphate buffered saline). CD3-CD19-CD56-CD14-HLA-DR+CD11c+CD1c cells were sorted. The average number of collected cells by sorting was 56,881 (range 6,669-243,385). For the second cohort, fresh PBMCs were immediately subjected to magnetic-activated cell sorting (MACS) for the removal (positive selection) of CD304+ cells (pDC), followed by the removal of CD19+ cells (B cell), and subsequently isolation of CD1c+ cells by using the CD1c+ (BDCA1) isolation kit (Miltenyi Biotec, Germany) according to the manufacturer’s instructions. The isolated CD1c+ fraction contained on average 147,114 cells (range 46,000-773,000) and purity was determined by flow cytometry (**Supplementary File 1B**) measured on the BD LSRFortessa Cell analyzer (**Figure 1 – Figure Supplement 1**). Data were analyzed using FlowJo software (TreeStar Inc.). MACS or FACS purified CD1c+ cells were immediately taken up in a lysis buffer (RLT plus, Qiagen) containing 1% β-mercaptoethanol, snap frozen on dry ice, and stored at −80°C until RNA extraction was performed. Isolation of CD1c+ DC for functional experiments was done by MACS as described above. Purification of CD1c+ DC subsets based on CD36 and CX3CR1 or CD14 expression from freshly isolated PBMCs was conducted by flow cytometry using the panel in **Supplementary File 1C** and shown in **Figure 3 – Figure Supplement 2B**.

### CD1c+ DC cultures and secretome analysis

Purified CD1c+ DCs were cultured in RPMI Glutamax (Thermo Fisher Scientific) supplemented with 10% heat-inactivated fetal bovine serum (FBS) (Biowest Riverside) and 1% penicillin/streptomycin (Thermo Fisher Scientific). CD1c+ DCs were cultured at a concentration of 0.5 × 10^6^ cells/mL in a 96-well round-bottom plate (100μL/well). Cells were stimulated overnight (18 hours) with multiple stimuli listed in **Supplementary File 1D**. After stimulation, cells were lysed in an *RLT plus* lysis buffer (Qiagen) and stored at −80°C until RNA extraction was performed. Cell lysates were stored at −80°C until RNA extraction was performed for qPCR. In separate cultures, CD1c+ DC subsets (sorted based on CD36 and CX3CR1 expression) were cultured in the presence of 1µg/mL lipoteichoic acid (LTA). After 18 hours of stimulation, supernatants were harvested and IL-23 cytokine production was analyzed by ELISA (R&D Systems). The levels of IL-2, IL-5, IL-6, IL-10, IL-12p70, IL-13, IL-17, IL-22, IL-27, TNF-alpha, IFN-alpha, IFN-beta, CCL1, CXCL10, CXCL13, VEGF, CD40L, FAS, TNFR1, TNFR2, Elastase, and Granzyme B were simultaneously measured in supernatant of CD1c+ DC cultures using the in-house multiplex immunoassay based on Luminex technology, as described previously (*26*). Protein concentrations that were out of range were replaced with the LLOQ (lower limit of quantification) and ULOQ (upper limit of quantification) for each analyte and divided by 2 for the proteins detected below the range of detection or multiplied by 2 for values above the detection range (**Supplementary File 1E**).

### Real-time Quantitative PCR

First-strand cDNA was synthesized from total RNA using Superscript IV kit (Thermo Fisher Scientific), and quantitative real-time PCR (RT-qPCR) was performed on the QuantStudio 12k flex System (LifeTechnologies), following manufacturer’s instructions. Sequences of the primers used are listed in **Supplementary File 1F** and the *Key Resource Table*. RT-qPCR data were normalized to the expression of the selected housekeeping gene *GUSB* (ENSG00000169919). CT values were normalized to GUSB by subtracting the CT mean of GUSB (measured in duplo) from the CT mean of the target mRNA (e.g., *RUNX3*) = ΔCT. The fold change (FC) of each sample was calculated compared to ΔCt of the medium control using the formula FC = 2−ΔΔCt, where ΔΔCt = ΔCt sample—ΔCt reference.

### Flow cytometry of CD1c+ DC populations

PBMC samples from the two cohorts (HC = 11 samples; AU = 9 samples; IU = 6 samples; BU = 11 samples) were randomly selected and measured by flow cytometry in batches of 9 to 10 mixed samples per run, divided over 4 days. Per batch, 10 million PBMCs per sample were quickly thawed, washed with ice cold PBS and stained with the antibody panel depicted in **Supplementary File 1C**. PBMCs were incubated with Fixable Viability Dye eF780 (eBioscience) at room temperature for 10 minutes. Cells were then plated in V-bottomed plates (Greiner Bio-one), washed with PBS and incubated for 30 minutes at 4°C in the dark with Brilliant Stain Buffer (BD) and the fluorescently-conjugated antibodies. Next, the cells were washed and taken up in the FACS buffer. Flow cytometric analyses were performed on the BD FACSAria™ III sorter. Manual gating of data was done using *FlowJo* software (TreeStar inc. San Carlos, CA, USA). FlowSOM v1.18.0 analysis was done as described previously (*27*). Lineage-(negative for CD3/CD56/CD19) HLA-DR+ data were transformed using the *logicleTransform* function of the *flowCore* v1.52.1 R package, using default parameters (*28*). The SOM was trained for a 7×7 grid (49 clusters) with 2000 iterations. Consensus hierarchical clustering was used to annotate clusters, based on the *ConsensusClusterPlus* v1.50.0 R package (*29*). Principal component analysis (PCA) analysis was done on normalized expression data from flowSOM using the *factoextra* v 1.0.7.999 R package.

### RNA isolation and RNA sequencing

Total RNA from CD1c+ DC cell lysates from patients and controls was isolated using the AllPrep Universal Kit (Qiagen) on the QIAcube (Qiagen) according to the manufacturer’s instructions. For cohort I, RNA-seq libraries were generated by *GenomeScan* (Leiden, the Netherlands) with the TruSeq RNAseq RNA Library Prep Kit (Illumina Inc., Ipswich, MA, USA), and were sequenced using Illumina HiSeq 4000 generating ~20 million 150 bp paired end reads for each sample. Library preparation and Illumina sequencing was performed on samples of cohort II at BGI (Hong Kong). RNA-seq libraries were generated with the TruSeq RNAseq RNA Library Prep Kit (Illumina Inc., Ipswich, MA, USA) and were sequenced using Illumina NextSeq 500 generating approximately 20 million 100bp paired end reads for each sample.

### Power analysis

We conducted power analysis using the PROPER R package v 1.22.0 (*30*) with 100 simulations of the *build-in* RNA-seq count data from antigen presenting (B) cells from a cohort of 41 individuals (i.e., large biological variation as expected in our study) (*31*). Simulation parameters used the default of 20,000 genes and an estimated 10% of genes being differentially expressed. We detected 0.8 power to detect differentially expressed genes (*P*<0.05) at a log_2_(fold change)>1 for the smallest patient group (9 cases) and we considered the sample size reasonable for analysis.

### Differential gene expression and statistical analysis

Quality check of the raw sequences was performed using the FastQC tool. Reads were aligned to the human genome (GRCh38 build 79) using STAR aligner (*32*) and the Python package HTSeq v0.6.1 was used to count the number of reads overlapping each annotated gene (*33*). We aligned the reads of the RNA sequencing data sets to 65,217 annotated *Ensemble Gene* IDs. Raw count data were fed into *DESeq2* v1.30.1(*34*) to identify differentially expressed genes (DEGs) between the four disease groups (AU, IU, BU, HC). Using DESeq2, we modeled the biological variability and overdispersion in expression data following a negative binomial distribution. We used Wald’s test in each disease group versus control pair-wise comparison and *P* values were corrected by the DESeq2 package using the Benjamini-Hochberg Procedure. We constructed co-expression gene networks with the WGCNA v 1.70-3 R package (*35*) using the cumulative uveitis-associated genes from all pairwise comparisons and a soft power of 5. Module membership (MM) represents the intramodular connectivity of genes in a gene module. Gene Significance (GS>0.25) indicates a strong correlation between genes and non-infectious uveitis, whereas MM (MM>0.8) indicates a strong correlation with the *EigenGene* value of the modules. We calculated the intersection between the modules constructed from the two cohorts and used Fisher’s exact test to identify modules that exhibited significant overlap in genes. Gene expression data from *runx3*-knockout(KO) cDC2s, *notch2*-KO cDC2s, and “inflammatory” cDC2s were obtained from the NCBI Gene Expression Omnibus (accession numbers GSE48590 [2 wild-type [WT] CD11b+ESAM+ splenic cDC2s versus 2 CD11b+ESAM+ cDC2s from CD11c-DC-*Runx3*_Δ_ mice], GSE119242 [2 untreated cDC2 versus untreated cDC2 from CD11c-Cre *notch2*f/f mice], GSE149619 [5 CD172+MAR1-cDC2s in mock condition vs 3 CD172+MAR1+ cDC2 in virus condition]) using GEO2R in the GEO database, which builds on the GEOquery v2.58.0 and limma R v 3.46 packages (*36, 37*). RNA-seq data from the mouse bone marrow stromal cell line OP9 expressing NOTCH ligand DLL1 (OP9-DLL1)-driven cDC2 cultures (GSE110577, [2 sorted CD11c+MHCII+B220−CD11b+ cDC2 from bone marrow cultures with FLT3L for 7 days vs 2 sorted CD11c+MHCII+B220−CD11b+ cDC2 from bone marrow cultures with FLT3L + OP9-DLL1 cells for 7 days]) were analyzed using DESeq2 and normalized count data plotted using the *plotCounts* function. RNA-seq count data from CD14+/-DC3 subsets from patients with SLE and systemic sclerosis were obtained via GEO (accession number: GSE136731) (*17*) and differential expression analysis was conducted using *DESeq2* v1.30.1. Gene set enrichment analysis was done using the *fgsea* R package v1.16.0 and data plotted using the *GSEA*.*barplot* function from the *PPInfer* v 1.16.0 R package (*38*). Gene sets for *runx3*-KO, *notch2*-KO, inflammatory cDC2s, and cDC2s from OP9-DLL1 bone marrow cultures were generated by taking the top or bottom percentiles of ranked [−log10(P) ⍰ sign(log2(fold change)] genes from each data set as indicated. Genes in the modules of interest that encode cell-surface proteins were identified according to *surfaceome* predictor *SURFY* (*39*);

### Single cell-RNA seq analysis of aqueous humor

Single cell RNA-seq (scRNA-seq) data from aqueous humor of 4 HLA-B27-positive anterior uveitis (identical to the AU group in this study) patients were obtained via Gene Expression Omnibus (GEO) repository with the accession code GSE178833. Data was processed using the R package *Seurat* v4.1.0 (*40*) using R v4.0.3. We removed low-quality cells (<200 or >2500 genes and mitochondrial percentages <5%) and normalized the data using the *SCTransform()* function accounting for mitochondrial percentage and cell cycle score (*41*). Dimensionality reduction for all cells was achieved by adapting the original UMAP coordinates for each barcode as reported by *Kasper et al 2021* (see GSE178833). Data were subjected to *scGate* v1.0.0 (*42*) using *CLEC10A+ and C5AR1* (CD88)- cells in our gating model to purify CD1c+ DCs in the scRNA seq dataset. Dimensionality reduction for CD1c+ DCs was conducted using the R package Seurat, and cells clustered using the *FindNeighbors* and *FindClusters* functions from Seurat. After clustering and visualization with UMAP, we used the *DotPlot* function from the Seurat package to visualize the average expression of genes in each cluster.

### Data and Code Availability

The data code (R markdown), bulk RNA-Seq datasets, flow cytometry dataset, and cytokine expression dataset described in this publication are available via https://dataverse.nl/doi: https://doi.org/10.34894/9Q0FVO and deposited in NCBI’s Gene Expression Omnibus accessible through GEO Series accession numbers GSE195501 and GSE194060.

## Results

### A CX3CR1 gene module is associated with non-infectious uveitis

We characterized the transcriptome of primary CD1c+ DCs from patients with non-infectious uveitis (**Figure 1A**). RNA-seq analysis (RNA-seq) was performed on lineage (CD3-CD19-CD56-CD14-)-negative, and HLA-DR-positive, CD11c and CD1c-positive DCs purified from frozen PBMCs by flow cytometry from 36 patients with anterior (AU), intermediate (IU), or posterior non-infectious uveitis (BU) and healthy controls. A co-expression network was constructed using uveitis-associated genes identified by differential expression analyses (n = 2,016 genes at *P<*0.05, **Figure 1B**), which identified six modules, of which 3 were associated with non-infectious uveitis (*Gene Significance* >0.25, **Figure 1C**). The blue module was most associated with non-infectious uveitis (**Figure 1C**). Based on *Module Membership, CX3CR1* was the top hub gene of the blue module (**Figure 1D, Supplementary File 1G**). Since CX3CR1 was previously associated with a distinct subset cDC2s that may also express CD14 (*43, 44*), we attempted to validate and expand the gene set associated with non-infectious uveitis by MACS-isolating CD1c+ DC cells from fresh blood of 28 patients and 14 healthy controls, followed by RNA-seq analysis of the highly purified CD1c+ DCs (median [interquartile range]% = 96[3]% pure, **Figure 1 - Figure Supplement 1**). We also constructed a co-expression network for uveitis-associated genes (n= 6,794, *P*<0.05) in the second cohort (**Figure 1E**), which revealed 24 gene modules (**Supplementary File 1H**). Note that patient samples did not cluster according to clinical parameters of disease activity (e.g., cell grade in eye fluid, macular thickness) (**Figure 1 - Figure Supplement 2**). The 3 uveitis-associated modules in cohort I shared a significant number of co-expressed genes with one module in cohort II, the black module (**Figure 1F**). The black module was associated with non-infectious uveitis in cohort II (Gene Significance for uveitis >0.25) and *CX3CR1* was also the hub gene for this module (Black Module Membership, *P* = 5.9 × 10^−22^; **Supplementary File 1H, Figure 1G**). According to these findings, the overlapping disease-associated gene modules appear to represent a single gene module. In cohort I, the separation of genes into three modules was possibly due to low sensitivity to detect disease-associated genes with low expression, as replicated genes of the black module were typically higher expressed (**Figure 1 - Figure Supplement 3**). In total, we replicated 147 co-expressed genes between the two cohorts (which we will refer to as the “black module”), of which 94% also showed consistent direction of effect (e.g., upregulated in both cohorts) (**Figure 1H, Supplementary File 1I**). The black module was enriched for the GO term “*positive regulation of cytokine production*” (GO:0001819, *Padj* = 6.9 × 10^−5^). In addition to *CX3CR1*, the black module comprised *CD36, CCR2, TLR-6, -7, -8, CD180*, and transcription factors *RUNX3, IRF8*, and *NFKB1* (**Figure 2I**), but not *CD14*. In summary, these results show that a gene module characterized by *CX3CR1* in blood CD1c+ DCs is associated with non-infectious uveitis.

**Figure 1.**
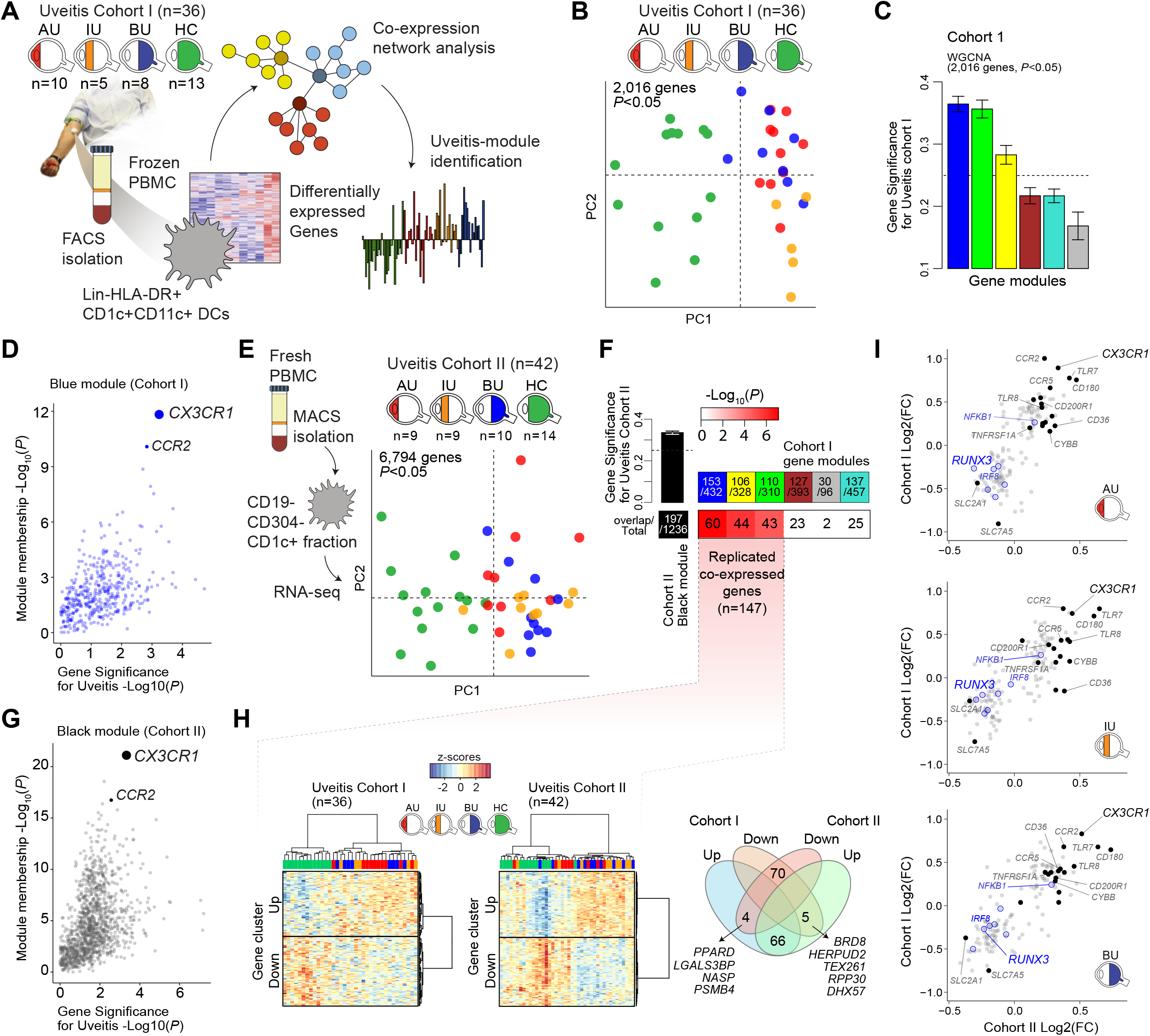
A *CX3CR1* gene module in CD1c+ Dendritic cells (CD1c+ DC) is associated with non-infectious uveitis. **A**) Study design. CD1c+ DCs were purified from blood and subjected to RNA sequencing. Co-expression network analysis was used to identify gene modules associated with uveitis. **B**) Principal component analysis (PCA) of the 2,016 uveitis-associated genes (*P*<0.05) in 36 patients and control samples of cohort I **C**) Gene significance for uveitis for the gene modules identified by WGCNA. **D**) Module membership and Gene Significance for uveitis for the blue module of cohort I. **E**) PCA of the 6,794 uveitis-associated genes (*P*<0.05) in 42 samples of cohort II. **F**) Cross-tabulation of the preservation of co-expressed genes between gene modules from cohort I and the black module from cohort II. *P* value is from Fisher’s exact test. **G**) Same as in *D*, but for the black module of cohort II. **H**) Heatmaps of the 147 replicated co-expressed genes (rows) for samples (columns) from cohort I and II. The venn diagram shows the upregulated and downregulated genes (clusters shown in *H*). **I**) The (Log2) fold change in gene expression compared to healthy controls (x-axis) for all 147 replicated genes in patients with AU, IU, AND BU. Genes encoding surface proteins are indicated in black/grey. Key transcription factors are indicated in blue. AU; Anterior uveitis. IU; Intermediate uveitis. BU; Birdshot uveitis.

**Figure 2.**
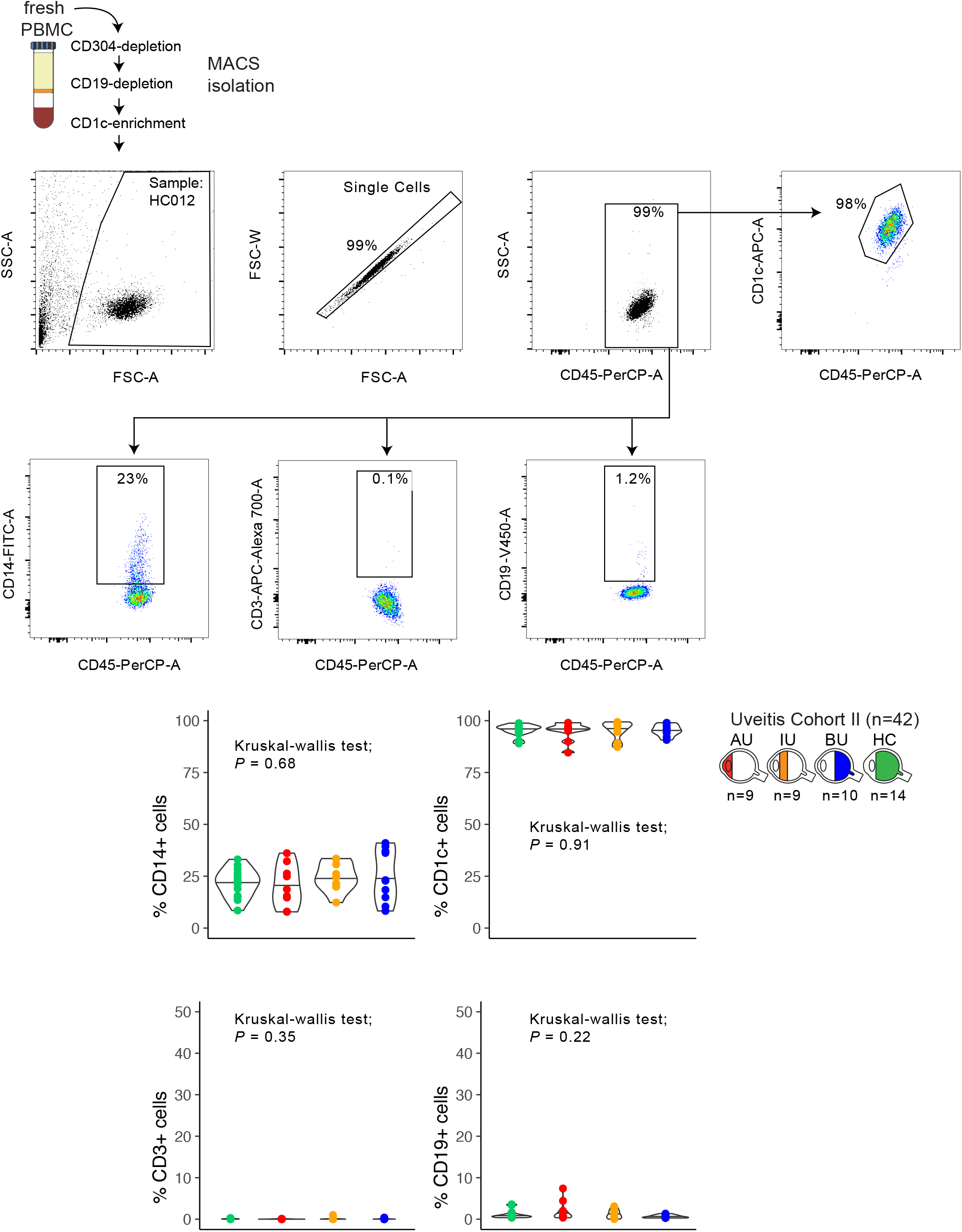
The *CX3CR1* gene module of CD1c+ DCs is enriched for NOTCH2-RUNX3 signaling. **A**) Volcano plot for the expression of genes of the black module in cDC2s of *runx3*-KO mice (GSE48590), *notch2*-KO mice (GSE119242), and type I IFN-dependent inflammatory [inf-]cDC2s (GSE149619). Up-regulated genes and down-regulated genes for each condition are indicated for each condition; grey dots denote the genes with no significant change in expression. **B**) Results from gene-set enrichment analysis for ranked transcriptomes (using 20,668 genes with baseMean>4) for AU, IU, and BU patients. The top or bottom percentiles of the ranked [−log10(P)l1l sign(log2(FC))] genes from runx3-KO cDC2s, notch2-KO cDC2s, and inf-cDC2s (see a) were used as gene sets. Normalized enrichment scores (NES) and *P* values for each gene set is indicated. The dotted lines indicate *Padj* = 0.05. AU = Anterior uveitis. IU = Intermediate uveitis. BU = Birdshot uveitis.

### CX3CR1+ DC3 are diminished in peripheral blood of non-infectious uveitis patients

Type I IFN cytokines promote differentiation of CD1c+ DCs (*17,19*), but patients with active non-infectious uveitis have reduced blood levels of type I IFN cytokines (*21,, 22*). Assessment of the transcriptome of CD1c+ DCs from patients, found no enrichment for genes associated with murine type I IFN-dependent cDC2s (*45*) (**Figure 2A-B**). Furthermore, in CD1c+ DCs from healthy human donors, IFN-alpha did not induce downregulation of *RUNX3* as observed in CD1c+ DCs from non-infectious uveitis patients (**Figure 2 - Figure Supplement 1A**). In contrast, the transcriptome of CD1c+ DCs from patients overlapped significantly with murine cDC2s knocked out for *Runx3*, or its upstream regulator *Notch2* (**Figure 2, Figure 2 - Figure Supplement 1B-C**) *(46–48)*. Given that cDC2 subsets differ by their dependence on NOTCH signaling (*46–49*), we hypothesized that the transcriptomic signatures of the CD1c+ DC pool in patients might reflect changes in their proportions.

Therefore, we used flow cytometry analysis to identify CD1c+ DC clusters in peripheral blood mononuclear cells (PBMCs) samples from 26 cases and 11 controls. We designed a panel based on the black module (CX3CR1, CD36, CCR2, and CD180), other CD1c+ DC markers that were *not* in the black module (CD1c, CD11c, CD14, CD5, and CD163) (*17, 50*). FlowSOM (*51*) was used on HLA-DR+ and lineage (CD3/CD19/CD56)-PBMCs to cluster cells into a predetermined number of 49 clusters (7×7 grid) to facilitate detection of CD1c+ DC phenotypes in blood. The analysis with flowSOM clearly distinguished four CD1c+ DC clusters (cluster number 22, 37, 44, and 45) (**Figure 3A** and **Figure 3 – Supplement 1A and 1B**). We extracted the data for these four CD1c+ DC clusters and conducted *principal component analysis* (PCA). The PCA biplot identified CD5 and CD163 as top loadings (**Figure 3 – Supplement 1C**), which defines the *DC2s* (cluster 45), CD5-CD163-DC3s (cluster 37), and CD5-CD163+ DC3s (cluster 22, and 44) (*17*) (**Figure 3B, 3C**). Among the identified clusters, we detected a significant reduction in the frequency of cluster 44 in patients compared to controls (*Welch T-test, P* = 0.03, **Figure 3D and Figure 3 – Supplement 1D**). Clusters 44 as well as cluster 22 were CD36 and CD14 positive, which indicates these clusters may represent CD14+ DC3s in human blood (*17*). However, cluster 44 had relatively higher levels of CX3CR1 than cluster 22. (**Figure 3E and 3F**). This suggests that DC3s may be phenotypically bifurcated by CX3CR1 independently of CD14. This is supported by weak correlation between *CD14* and *CX3CR1* in our RNA-seq data from bulk CD1c+ DCs (Pearson correlation coefficient = 0.35, **Figure 3 – Supplement 2A**). In addition, we sorted CD14-positive and CD14-negative fractions from CD1c+ DCs of six healthy donors (**Figure 3 - Figure Supplement 2B**) which showed no significant difference in expression levels for CX3CR1 (**Figure 3 - Figure Supplement 2C**). *CX3CR1* levels were also not significantly different in sorted CD5-CD163+ CD14-positive and CD14-negative DC3s from patients with autoimmune diseases, further indicating that *CX3CR1* expression in CD1c+ DCs may be independent from *CD14* expression in CD1c+ DCs (**Figure 3 - Figure Supplement 2D**).

**Figure 3.**
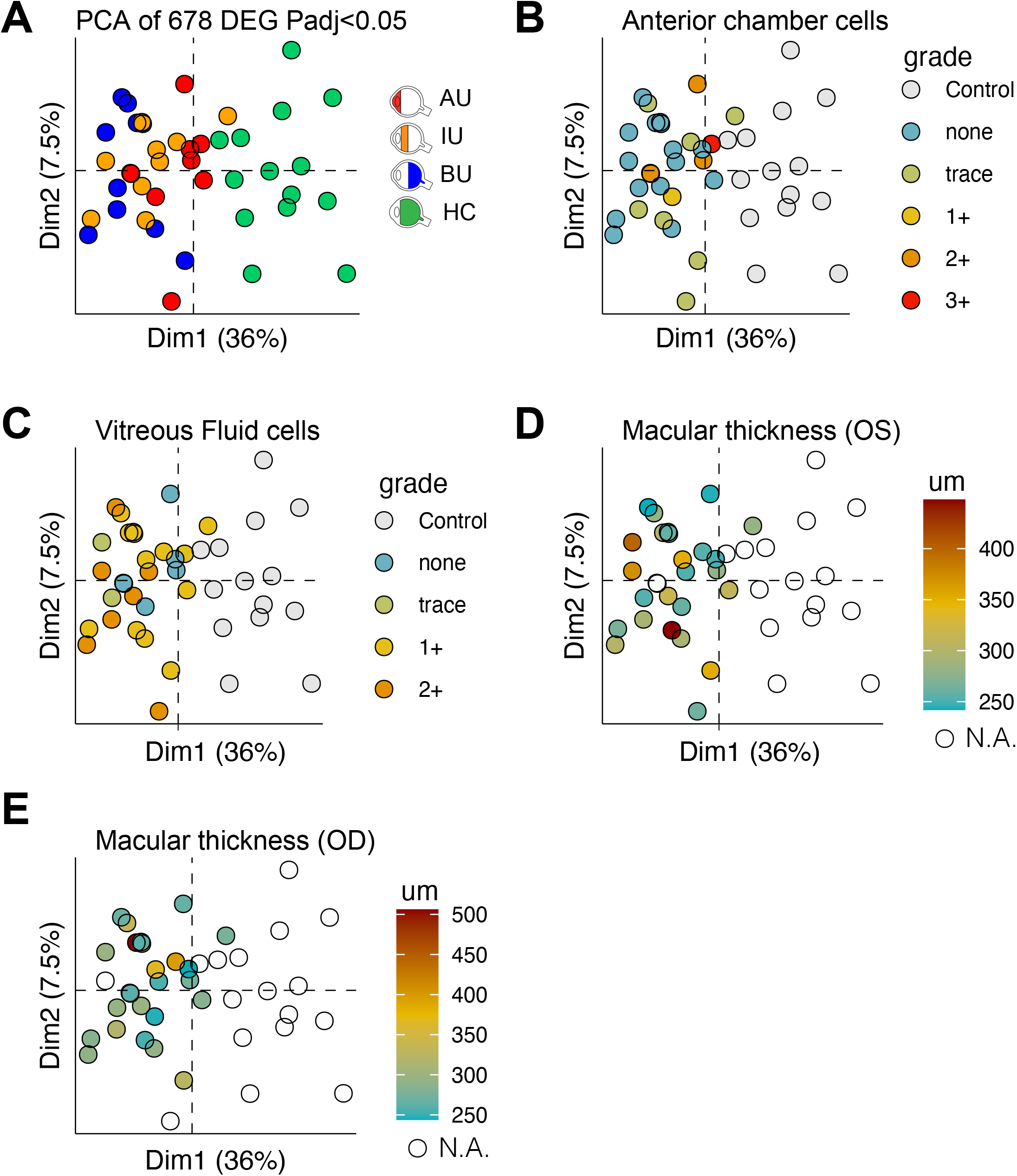
CX3CR1+ DC3s are decreased in the blood of patients with non-infectious uveitis. **A**) Heatmap of the surface protein expression for 49 flowSOM clusters of flow-cytometry analysis of PBMC samples from 26 patients and 11 controls. The four CD1c+ (CD3-CD19-CD56-HLA-DR+CD11c+) DC clusters identified (cluster 22, 37, 44, and 45) are shown (detailed heatmap in *Figure 3 – Supplement 1A*). **B**) Biplot of the normalized surface expression of CD5 and CD163 for the 4 CD1c+ DC clusters. **C**) Correlation plot between manually gated CD5-CD163-DC3s and CD5-CD163+ DC3s and DC3 flowSOM clusters 22, 37, and 44. **D**) The frequency of the 4 CD1c+DC flowSOM clusters as percentage of PBMCs. *P* values from *Welch’s t-test*. **E**) PCA biplot of the DC3 clusters 22, 37, and 44. Loadings for PC1 and PC2 are shown on the right. **F**) Biplots of the normalized surface expression of CD36, CD14, and CX3CR1 in the DC3 clusters 22, 37, and 44. **G**) Manual gating strategy of CD1c+ DC subsets based on CD36 and CX3CR1 in PBMCs in uveitis cases and controls. *P* value from *Welch’s t* test. Details on manual gating strategy: see *Figure 3 – Supplement 3*. Manual gating revealed that the CD14+ CD1c+ DCs (DC3s) can be further subdivided in a CX3CR1- and a CX3CR1+ population.

We validated by manual gating that cluster 44 represents the CD36+CX3CR1+ fraction of CD1c+ DCs in peripheral blood (~25% of total CD1c+ DCs) (**Figure 3 - Figure Supplement 1E**). Comparison between patients and controls corroborated that the frequency of manual gated CD36+CX3CR1+ DC3s were decreased in the blood of non-infectious uveitis patients (Welch *t-*test, *P* = 0.029, **Figure 3G**). In detail, we show that CD14+ CD1c+ DCs double positive for CD36+ and CX3CR1 were significantly decreased (*P* = 0.026), while CD14+CD1c+ DCs not positive for CX3CR1 were not (*P* = 0.43) (**Figure 3G**). This supports that CX3CR1 discerns a phenotypic subpopulation of CD14+ DC3s (**Figure 3 - Figure Supplement 3**), that was diminished in the blood of patients with non-infectious uveitis.

### CX3CR1+DC3s can secrete pro-inflammatory cytokines upon stimulation

We compared the cytokine-producing abilities of CX3CR1+ DC3s to their negative counterparts since the gene module associated with *CX3CR1* was enriched for genes involved in cytokine regulation. To this end, we freshly sorted primary human CD1c+ DC subsets based on the surface expression of CX3CR1 and CD36, of which double-positive and double-negative subsets could be sorted from the selected healthy subjects in sufficient numbers for analysis (**Figure 4 – Figure Supplement 1**). Since CD36 is involved in lipoteichoic acid (LTA) induced cytokine production (*52*), we overnight stimulated the CD1c+ subsets with LTA. Both subsets of CD1c+ DCs secreted IL-23 equally strongly (**Figure 4A**). To assess the secretome of the CD1c+ DC subsets in more detail, we profiled the supernatants of LTA-stimulated CD1c+ DC subsets for additional soluble immune mediators (**Supplementary File 1E**): The CD1c+ DC subsets could be distinguished based on the secreted protein profile (**Figure 4B**), of which the levels of TNF-alpha, IL-6, VEGF-A, and TNFR1 showed significant differences between the subsets (**Figure 4C**). These results show that CD1c+ DC subsets defined on the basis of surface co-expression of CD36 and CX3CR1 have the capacity to secrete pro-inflammatory mediators that participate in the pathophysiology of human non-infectious uveitis.

**Figure 4.**
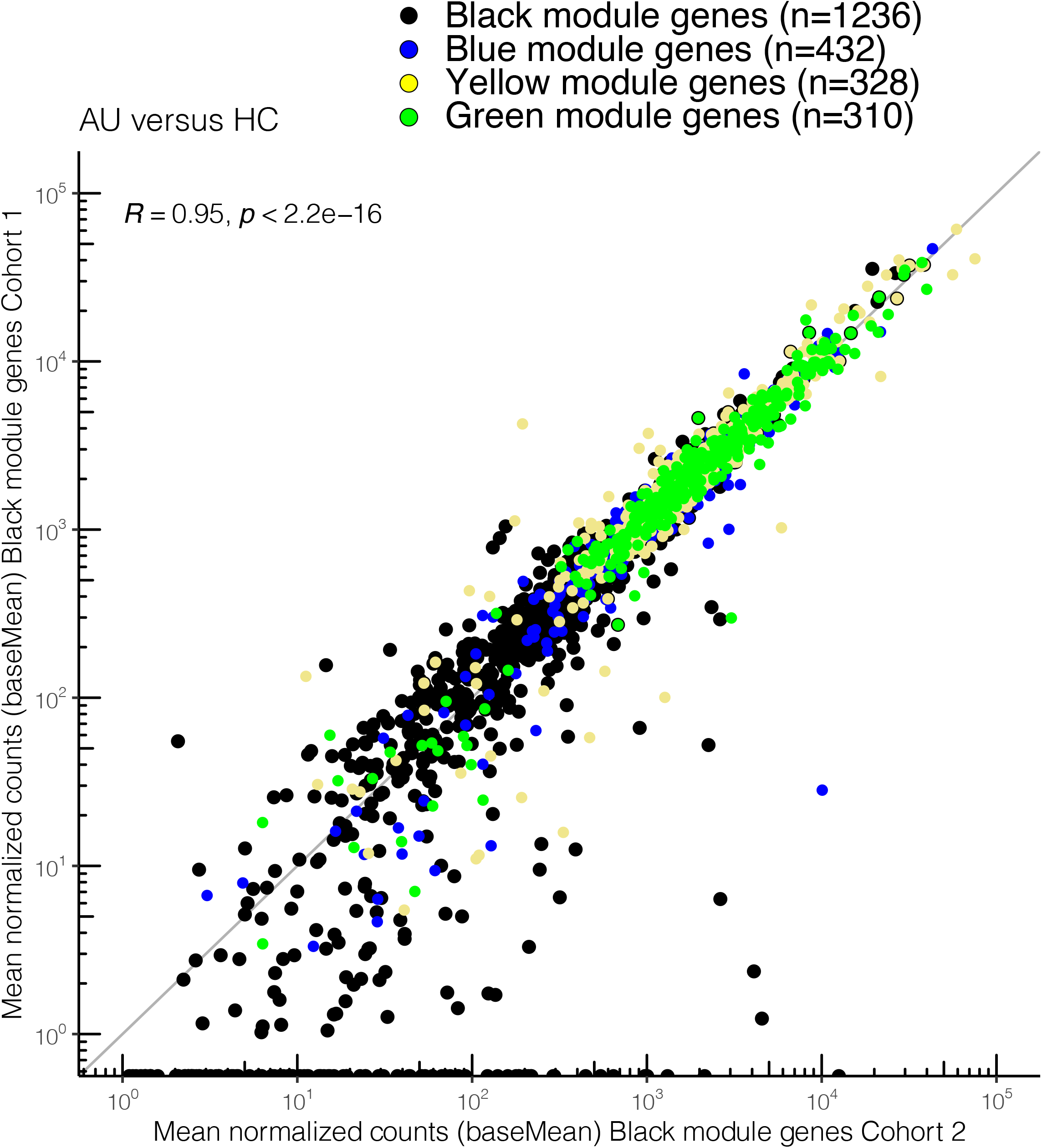
CX3CR1+DC3s secrete high levels of cytokines implicated in non-infectious uveitis. **A**) The CD1c+ DC cells were FACS sorted into CD36+CX3CR1+ and CD36-CX3CR1-CD1c+ DCs (Figure 4 – Supplement 1). The concentration of IL-23 (ELISA) in supernatants of 18h cultured primary human CD1c+ DC subsets cells stimulated with lipoteichoic acid (LTA). **B**) Heatmap of the levels (Z-score) of 16 detected proteins in supernatants of 18h cultured LTA-stimulated primary human CD1c+ DC subsets cells using an in-house multiplex *Luminex* assay (**Supplementary File 1E**). **C**) Scatter plots with overlay boxplot with mean and interquartile range of the levels of secreted TNF-alpha, Interleukin (IL)-6, VEGF-A, and TNFR1 from the multiplex protein data in *d*. (*Padj* = *P* values from likelihood ratio test Bonferroni corrected for 16 detected proteins).

### CX3CR1+ DC3s are detectable in the inflamed eye during non-infectious uveitis

We speculated that CX3CR1+ DC3s are important in the disease mechanisms of uveitis and may be found at increased abundance in the eye during active uveitis. We used single-cell RNA sequencing data (scRNA-seq) of eye fluid biopsies of 4 noninfectious patients (*53*). Cells positive for the CD1c+ DC specific tissue-marker *CLEC10A* and negative for the monocyte marker *C5AR1* (CD88) (*17, 20, 54*) were used to identify CD1c+ DCs among other immune cells in the scRNA-seq data (**Figure 5A**). Unsupervised clustering identified three clusters (1, 2, and 3) of different cells within the CD1c+ DC population (**Figure 5B)**. We identified that cluster 1 expressed the gene profile associated with CX3CR1+ DC3s, including relatively higher levels of *CX3CR1, CD36, CCR2*, and lower levels of *RUNX3* compared to the other two CD1c+ DC clusters (**Figure 5C**), which is in line with the gene profile identified by our bulk RNA-seq analysis. In summary, we conclude that CD1c+ DCs with a gene expression profile similar to CX3CR1+ DC3s can be detected in the eyes of patients during active non-infectious uveitis.

**Figure 5.**
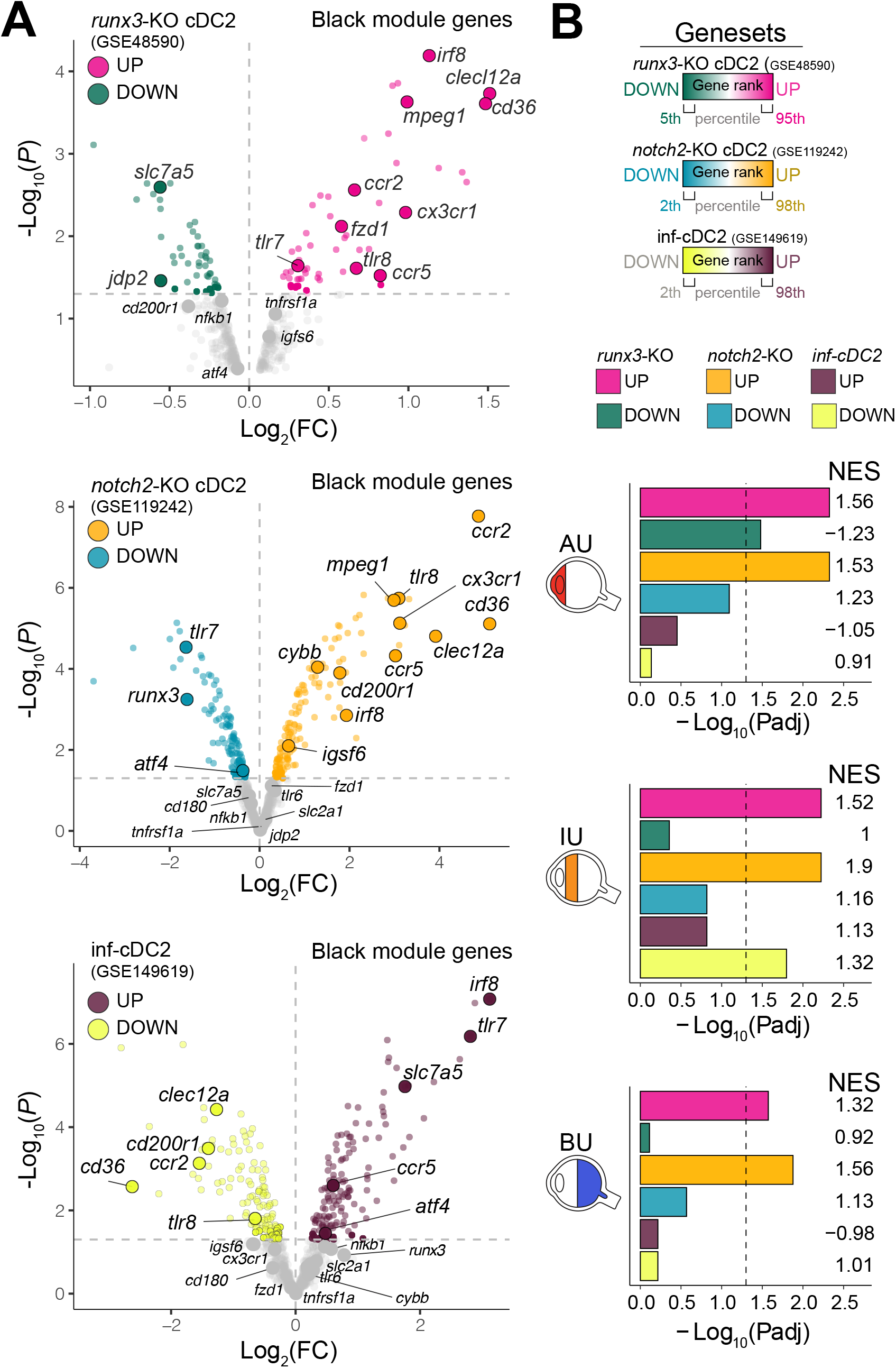
Local CD1c+ DCs show a gene profile similar to CX3CR1+ DC3s in the inflamed eye during non-infectious uveitis. **A**) Single-cell RNA-sequencing (scRNAseq) analysis of eye fluid biopsies from non-infectious uveitis patients (*GSE178833*). UMAP projections of transcriptomic data from 492 CD1c+ DC cells (in red) identified by *scGate* analysis (using *CLEC10A*+ and *C5AR1*-cells as tissue markers for CD1c+ DCs). **B**) Unsupervised clustering of CD1c+ DCs identified in *a*. **C**) Dot plot showing average expression (color-scaled) of key marker genes of the black module and *CD14* in each cluster determined in *B*.

## Discussion

In this study of non-infectious uveitis patients and controls, we identified and replicated a *CX3CR1*-associated gene module in CD1c+ DCs. We were able to track back the gene module to a CX3CR1+ DC3 subset that was diminished in peripheral blood of patients with non-infectious uveitis.

Preceding studies into human CD1c+ DCs revealed functionally distinct subsets termed “DC2” and “DC3”, with the DC3 showing both transcriptomic features reminiscent of cDC2s and monocytes - such as elevated *CD36* (*16,,17*). DC3s also have distinct developmental pathways and transcriptional regulators compared to DC2 (*16-18,20*). Recently, Cytlak and associates revealed that lower expression of *IRF8* is linked to DC3 (*18*), a transcription factor that was also decreased in non-infectious uveitis. According to Brown et al. (*43*), CD1c+ DCs exhibit two subsets: cDC2A and cDC2B, whereas cDC2B exhibits higher expression of CX3CR1 and produces more TNF alpha and IL-6 than cDC2A upon stimulation. Accordingly, uveitis-associated CX3CR1+ DC3s described in this study exhibit similar phenotypical and functional features.

*Dutertre* and co-workers (*17*) showed that the phenotype of peripheral blood CD1c+ DCs can be further segregated according to the expression of CD163 and CD5, with “DC3” cells being characterized as CD5-CD163-or CD5-CD163+cells and “DC2” as CD5+CD163 cells. Our flow cytometry results confirm these findings, but we also show that CD5-CD163+ DC3s that express CD14 are composed of CX3CR1-positive and CX3CR1-negative cells, of which the CX3CR1+ population is implicated in non-infectious uveitis.

Single-cell analysis supported that CD1c+ DCs in eye fluid of patients with noninfectious uveitis contain also a population that has a gene profile reminiscent of CX3CR1+ DC3s, with relatively higher levels of CX3CR1, CD36, CCR2, and lower levels of RUNX3. Patients with Systemic lupus erythematosus (SLE) display accumulation of CD14+DC3s in blood (*17*), while the population of CD14+DC3 cells was decreased in non-infectious uveitis patients. The differences between non-infectious uveitis and SLE may be related to distinct (i.e., opposite) immunopathological mechanisms; Type I interferons drive the maturation of cDC2s into “inflammatory cDC2s” (infcDC2s) (*45*) and can induce CD1c+ DCs to express a distinct set of surface-receptors (*19*). The type I interferon (IFN)-α drives immunopathology of SLE and administration of type I interferon therapy can induce *lupus-like* disease (*23,24*). In favor of attributing the seemingly contrasting observations in blood CD1c+ subsets between SLE and non-infectious uveitis to distinct biology is the fact that, in contrast to elevated IFN-α in patients with SLE, in non-infectious uveitis patient’s disease exacerbations correlate with reduced blood type I IFN concentrations (*21,22,55*). Despite the importance of type I IFN signaling on DC3s, our results suggest that DC3s are also dysregulated in conditions associated with decreased type I IFNs (*21,22*), supporting additional pathways involved in DC3 regulation during chronic inflammation.

We showed that the gene module of CD1c+ DCs showed overlap with the gene signature of disrupted NOTCH2-signaling in cDC2s. *Notch2* signaling is mediated via the NF-κB family member *Relb* in murine cDC2s (*56*). NF-κB signaling via RelB suppresses type I Interferon signaling in cDC2s (*57*) while selective deletion of RelB in dendritic cells protects against autoimmunity (*56,58*). It is tempting to speculate that the enrichment for NOTCH gene signatures implies altered NF-κB-Relb signaling in CD1c+ DCs, with a mechanism that varies between diseases mediated by type I IFNs (e.g., SLE) and type I IFN-negative diseases (e.g., uveitis). Although this warrants further investigation, some circumstantial evidence for this is the presence of NF-κB family members in the black module, such as *NFKB1* and *NFKBIA*, the former associated previously with a CX3CR1+ cDC2s, while the latter can regulate *Relb* function in dendritic cells (*59,60*). NF-κB-Relb signaling has been shown to suppress type I IFN via a histone demethylase encoded by *KDM4A* which was also in the black module (*61*). Regardless, the NF-κB pathways are regulated by the TNFR1, the main receptor for TNF alpha (*62, 63*). TNFR1 is expressed at the cell surface of cDC2s and its ectodomain cleaved by the NOTCH2-pathway regulator ADAM10 (63–65). We showed that both TNF alpha and sTNFR1 were higher in the secretome of activated CX3CR1+ DC3s. This is in agreement with previous studies on CD36+ DC3s (*16*) or CX3CR1+ cDC2B (T-bet–) that also produced higher levels of TNF-alpha (*43*). Interestingly, altered NF-κB signaling specifically in cDC2 is associated with clinical response to anti-TNF alpha therapy. (66). Anti-TNF therapy is effective for treatment of non-infectious uveitis (*67*), while anti-TNF therapy may also result in a dysregulated type I interferon response (*68*) indicating potentially cross regulatory mechanisms via NF-κB signaling and type I IFN signaling affecting cDC2s. More research is needed to resolve the regulatory mechanisms driving CD1+ DC changes in type I IFN-negative inflammation, including non-infectious uveitis. It is also possible that the change in CD1c+ DCs observed in this study results from cytokine-induced precursor emigration or differentiation.

Other disease modifying factors possibly affect the CD1c+ DC pool in uveitis patients. In mice, antibiotic treatment to experimentally disturb the microbiota affects a cDC2 subset phenotypically similar to CD1c+ DCs and decreases their frequency in the intestine of mice, which suggests microbiota-dependent signals involved in the maintenance of cDC2 subsets (*43*). This is especially interesting in light of growing evidence that microbiota dependent signals cause autoreactive T cells to trigger uveitis (*69*), which makes it tempting to speculate that gut-resident cDC2 subsets contribute to the activation of T cells in uveitis models. Dietary components can influence subsets of intestinal dendritic cells (*70*). Regardless, most likely, an ensemble of disease modulating factors is involved. For example, myeloid cytokines, such as GM-CSF, contribute to autoimmunity of the eye (*71*) and GM-CSF has been shown to stimulate the differentiation of human CD1c+ subset from progenitors (*20*). However, GM-CSF signaling in conventional dendritic cells has a minor role in the inception of EAU (*72*). Our data supports that stimulation of CD1c+ subsets with GM-CSF or TLR ligands does not induce the transcriptional features of CD1c+ DCs during non-infectious uveitis, which is in line with previous observations that support that stimulated cDC2s do not convert from one into another subset (*20*).

Note that our results of a decreased subset of CD1c+ DCs in non-infectious uveitis are in contrast with previous flow cytometry reports in non-infectious uveitis (12-14). Chen and co-workers (*14*) reported an increase of CD1c+ myeloid dendritic cells in non-infectious uveitis. It is important to note, however, that their study did not include the DC marker CD11c, thereby including CD1c+CD11c-populations that do not cluster phenotypically with CD1c+ (CD11c+) DCs (e.g., *cluster 46*, se **Figure 3 - Figure Supplement 1A**), which may explain the differences compared to our study.

Better understanding of the changes in the CD1c+ DC pool during human non-infectious uveitis will help develop strategies to pharmacologically influence putative disease pathways involved at an early disease stage, which may lay the foundation for the design of effective strategies to halt progress towards severe visual complications or blindness. Perhaps targeting CD1c+ DCs may be achieved by dietary (microbiome) strategies and provide relatively safe preventive strategies for noninfectious uveitis.

To conclude, we have found that peripheral blood CD1c+ DCs have a gene module linked to a CX3CR1-positive CD1c+ DC subset implicated in noninfectious uveitis.

## Supplementary Materials

### Figure Supplements

Supplementary File 1A-I

## Competing interests

Authors declare that they have no competing interests

## Figure Legends

**Figure 1 - Supplement 1.**
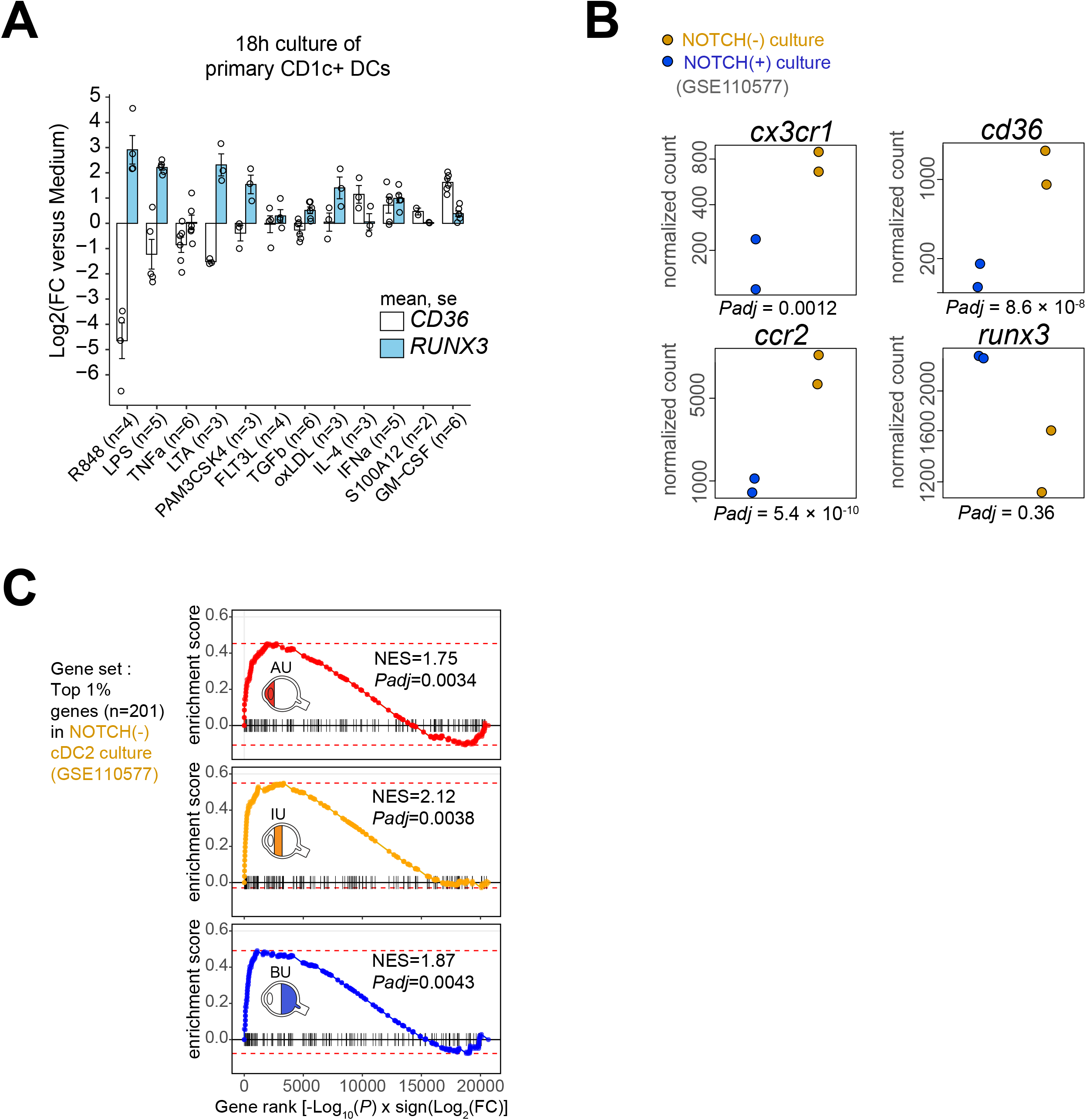
Purity check of cell fractions for RNA-sequencing in cohort I. Representative sample of flow cytometry gating of CD14+, CD19+, CD3+ and CD1c+ cell fractions in CD304-depleted, CD19-depleted and CD1c+ enriched MACS fractions from fresh peripheral blood mononuclear cells. Manual gating data for each individual sample is available via: https://doi.org/10.34894/9Q0FVO. The percentage of cells positive for each marker on the group levels between the disease groups is indicated in the bottom.

**Figure 1 - Supplement 2.**
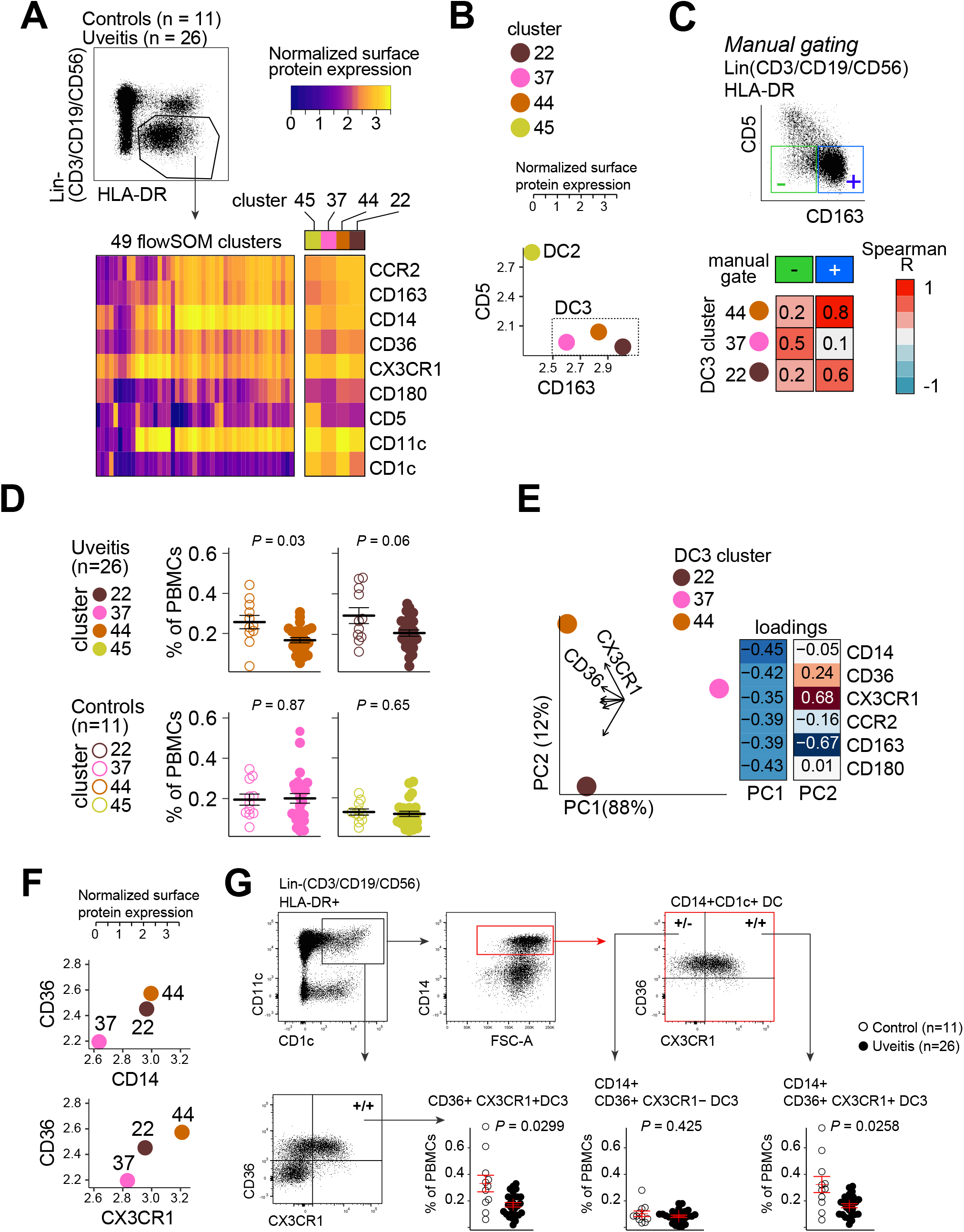
Clinical parameters of disease activity in non-infectious uveitis in cohort I. **A-E**) A PCA plot based on the 678 differentially expressed genes (*Padj*<0.05). (**A**) the anterior chamber cell grade (**B**), Vitreous fluid cell grade (**C**), macular thickness in the left (OS) eye as determined by optical coherence tomography (OCT) (**D**), and macular thickness in the right eye (OD) as determined by OCT (**E**) are shown.

**Figure 1 - Supplement 3.**
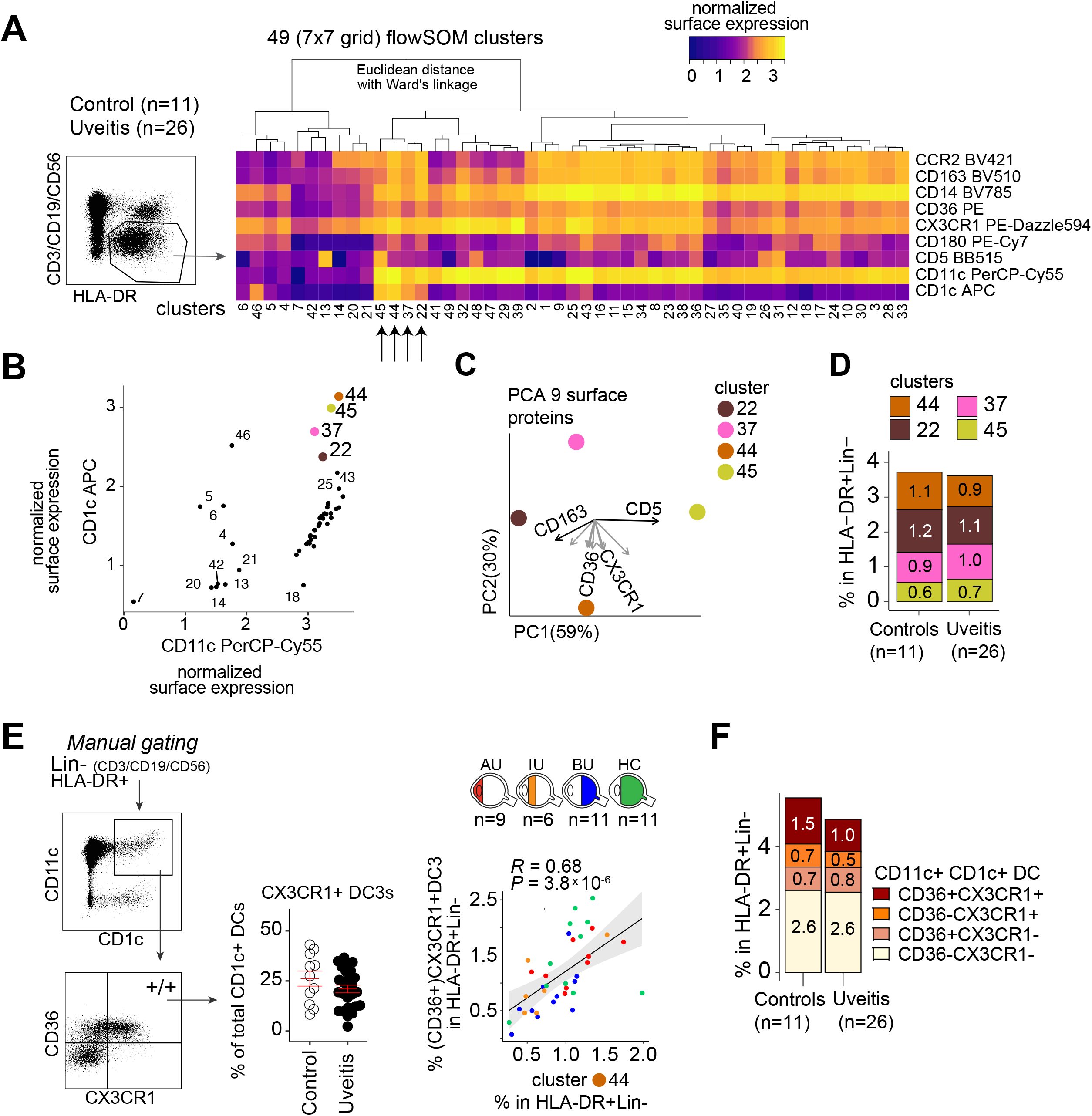
Correlation plot of the mean normalized count (*baseMean* from *DESeq2*) of the black module genes from cohort 1 and the 147 overlapping genes in the blue, yellow, and green module in cohort 2.

**Figure 2 - Supplement 1.**
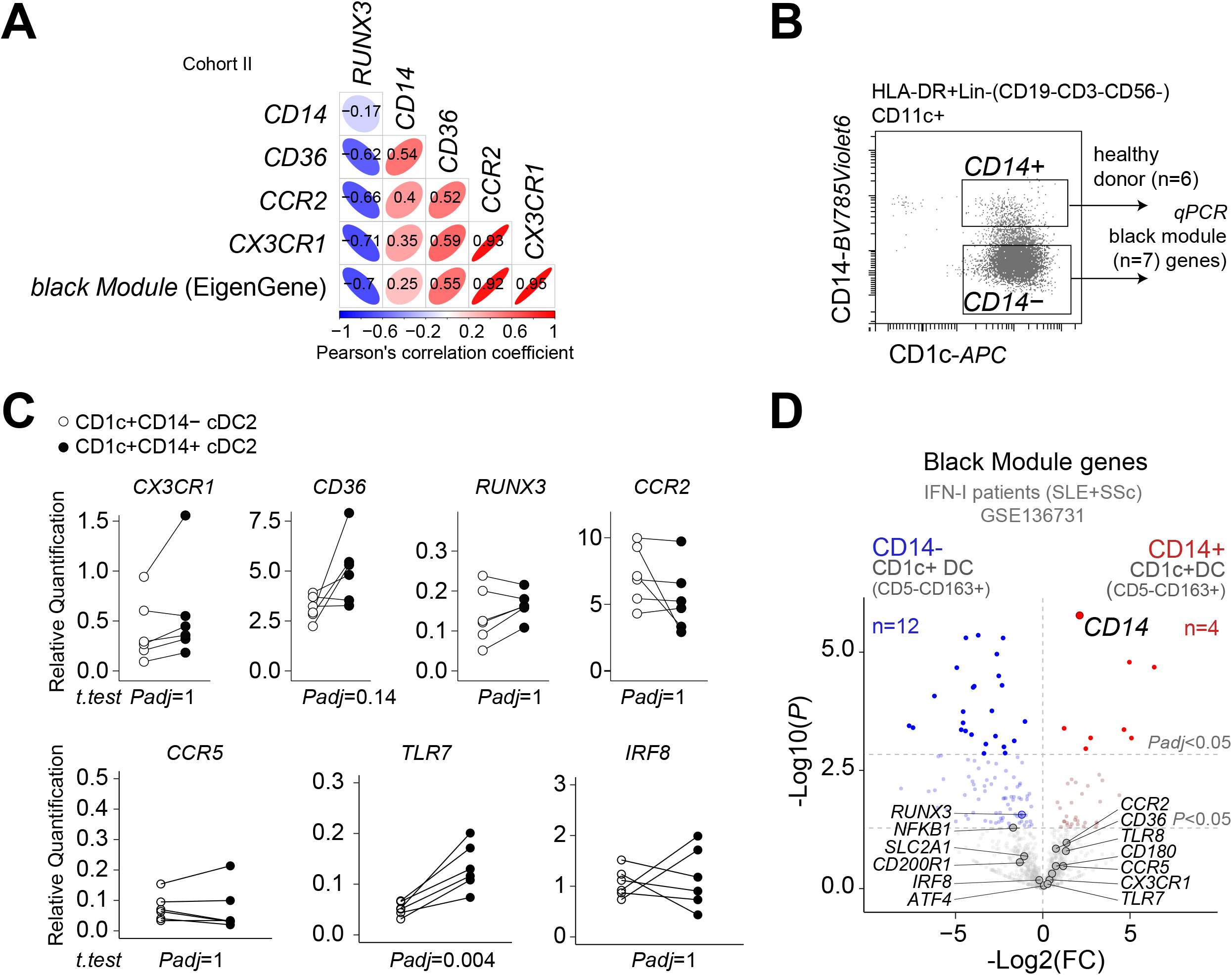
**A**) Gene expression (mean [SEM]) for *RUNX3* and *CD36* in primary human CD1c+ DCs from healthy donors stimulated overnight. Each dot represents a single donor used in the experiment. **B**) Transcriptomic data of murine bone marrow progenitors cultured for 7 days with OP9 stromal cells that express the NOTCH2 ligand DLL1 or OP-9 cells without DLL1 This analysis revealed that notch2-controlled genes were enriched in the transcriptome of CD1c+ DCs of patients and that notch2-signaling associates with the expression of *cd36, ccr2*, and *cx3cr1* in cDC2s. Normalized counts (and adjusted *P* values from *DESeq2*) for *cx3cr1, ccr2, cd36*, and *runx3* from cDC2s (GSE110577) generated from murine bone marrow cells and OP-9 with (in blue) or without (in ochre) Notch ligand Delta-like 1 (DLL1). **C**) Gene set enrichment analysis using the top 1%, (n=201) genes associated with the NOTCH-negative condition in *d* as the gene setR848 = Resiquimod, LTA = Lipoteichoic acid, LPS = Lipopolysaccharides, Pam3CSK4 = Pam3CysSerLys4, OxLDL = Oxidized low-density lipoprotein, IFNα = Interferon alpha, TGFβ = transforming growth factor beta. FLT3L = FMS-like tyrosine kinase 3 ligand, TNFα = tumor necrosis factor alpha, S100A12 = S100 calcium-binding protein A12, IL-4 = interleukin 4, GM-CSF = Granulocyte-macrophage colony-stimulating factor.

**Figure 3 - Supplement 1.**
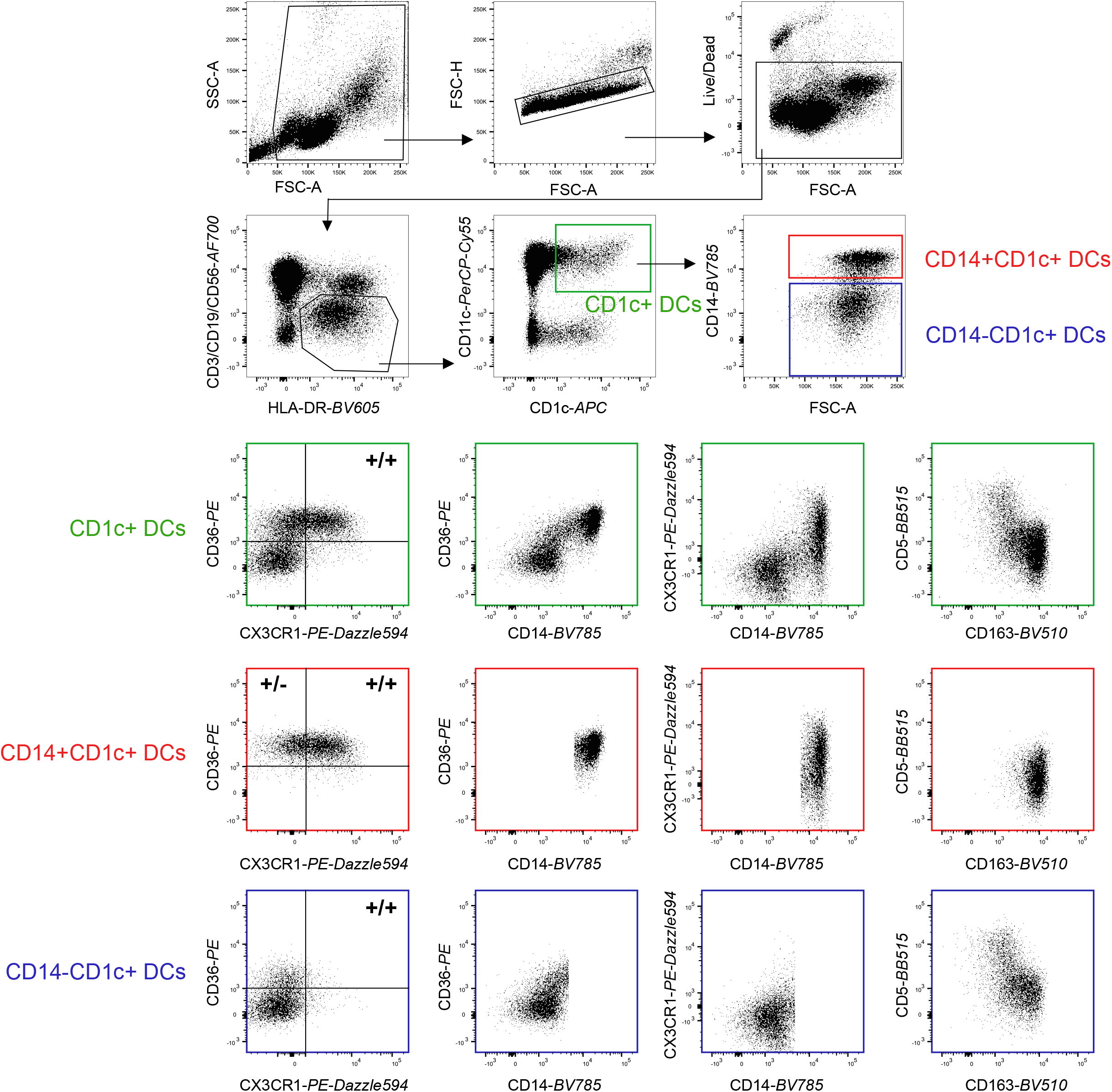
**A**) Heatmap of the surface protein expression for 49 flowSOM clusters of flow-cytometry analysis of PBMC samples from 26 patients and 11 controls. The four CD1c+ (CD3-CD19-CD56-HLA-DR+CD11c+) DC clusters identified (cluster 22, 37, 44, and 45) are highlighted. **B**) Biplot of the cell surface expression of CD1c and CD11c for the 49 flowSOM clusters in *a*. **C**) PCA biplot of the surface protein expression for clusters 22, 37, 44, and 45 identified in *a*. **D**) The proportion of the 4 CD1c+ DC clusters in the HLA-DR+Lin-(CD3-CD19-CD56) population in controls and patients. **E**) Manual gating strategy of CD1c+ DC subsets based on CD36 and CX3CR1 and the correlation between CD36+CX3CR1+ CD1c+ DCs and the flowSOM cluster 44. *R* = Spearman correlation. Gray area represents the 95% confidence interval of the linear regression line. **H**) The relative proportion of CD1c+ DC subsets based on CD36 and CX3CR1 in the HLA-DR+ Lin-gate.

**Figure 3 - Supplement 2.**
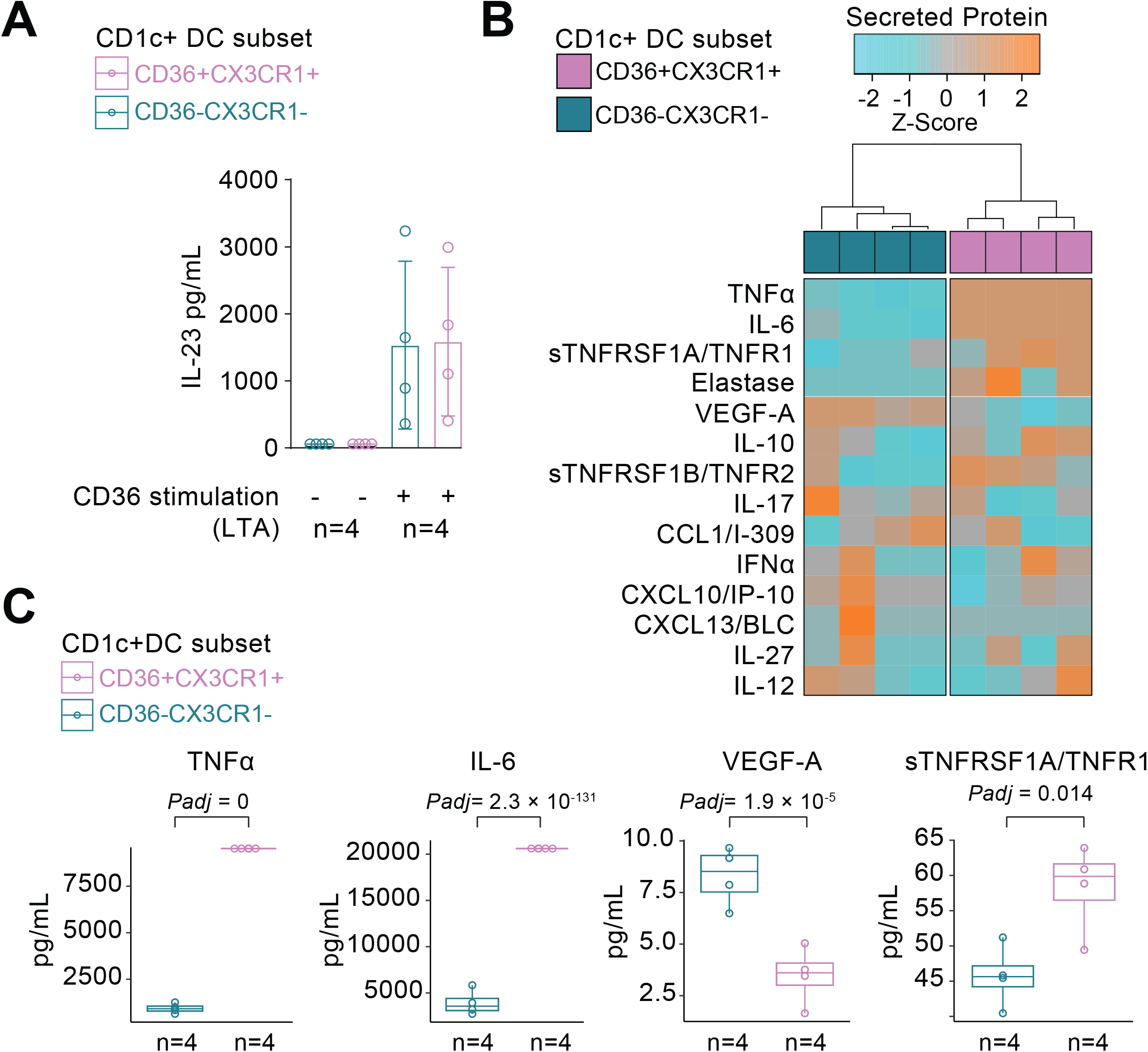
**A**) Correlation plot of the gene expression levels of *CD14, CD36, CCR2, CX3CR1*, and the *EigenGene* value of the black module. **B**) Gating strategy to sort CD14+ and CD14-CD1c+ DCs from healthy donors. **C**) RT-qPCR results for a panel of genes of the black module in the sorted CD14+ and CD14-CD1c+ DC fractions in *a. Padj* = adjusted *P* values from t.test (Bonferroni) corrected for 7 genes. **D**) Volcano plot showing differential expression analysis for black module genes in CD14+ versus CD14-DC3s purified from SLE and SSc patients (GSE136731).

**Figure 3 - Supplement 3.**
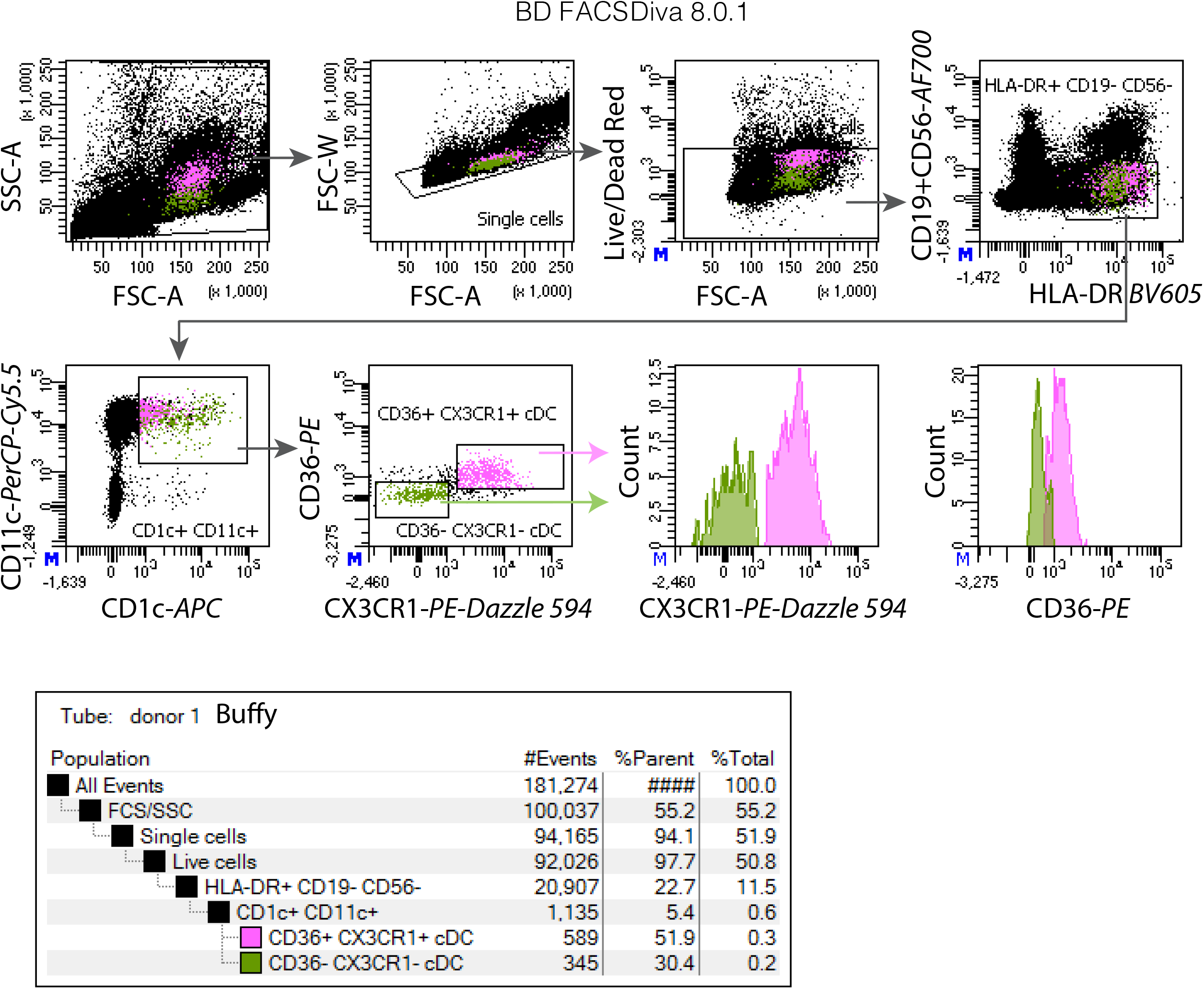
Representative sample of flow cytometry gating of CD14+ and CD14-fractions of CD1c+ DCs in peripheral blood for the panel used in Figure 3.

**Figure 4 - Supplement 1.**
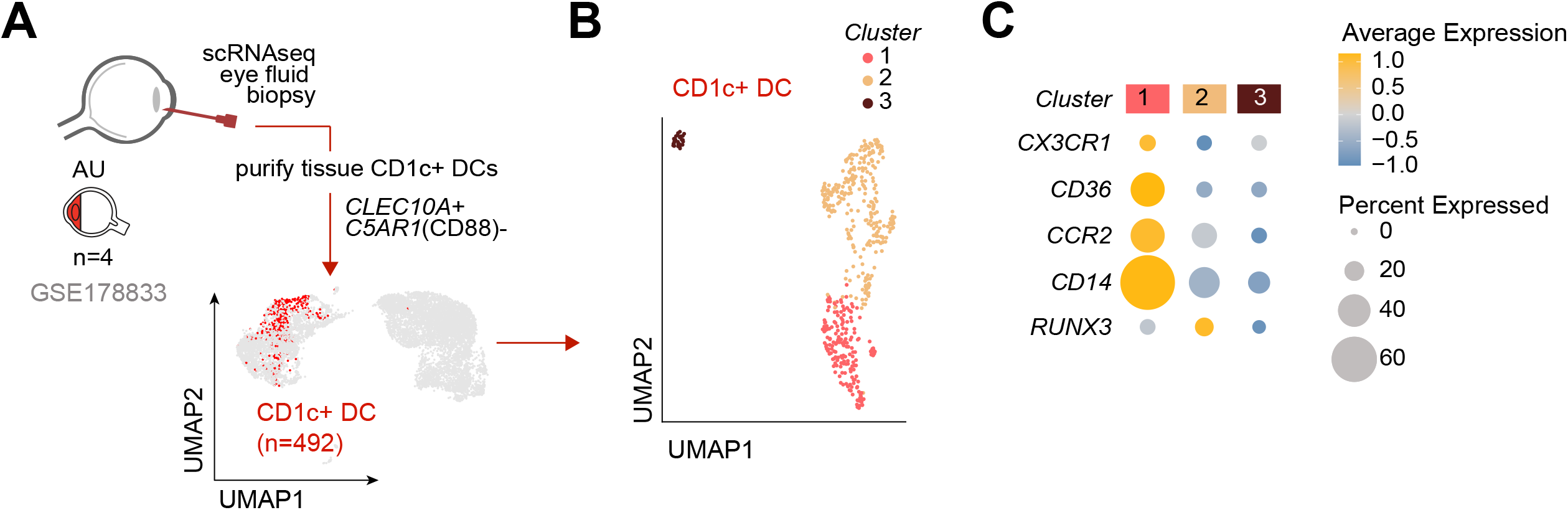
Representative examples of fluorescent-activated cell sorting (FACS) of CD36+CX3CR1+ DC3s and CD36-CX3CR1-CD1c+ DCs used for the analysis in Figure 4.

## Notes

### Competing Interest Statement

The authors have declared no competing interest.

### Summary of Updates

Revision after reviewer comments.

